# Efficient formation and maintenance of humoral and CD4 T cell immunity targeting the viral capsid in acute-resolving hepatitis E infection

**DOI:** 10.1101/2023.10.19.563038

**Authors:** Benedikt Csernalabics, Mircea Stefan Marinescu, Lars Maurer, Lara Kelsch, Jill Werner, Katharina Baumann, Katharina Zoldan, Marcus Panning, Philipp Reuken, Tony Bruns, Bertram Bengsch, Christoph Neumann-Haefelin, Maike Hofmann, Robert Thimme, Viet Loan Dao Thi, Tobias Boettler

**Affiliations:** Department of Medicine II, Medical Center – University of Freiburg, Germany; Faculty of Medicine, University of Freiburg, Germany; Schaller Research Group, Department of Infectious Diseases and Virology, Heidelberg University Hospital, Germany; Institute of Virology, University Hospital Freiburg, Germany; Department of Internal Medicine IV, University Hospital Jena, Germany; Department of Internal Medicine III, University Hospital RWTH Aachen, Germany; Signalling Research Centres BIOSS and CIBSS, University of Freiburg, Freiburg, Germany; German Centre for Infection Research (DZIF), Partner Site Heidelberg, Heidelberg, Germany

**Keywords:** adaptive immunity, immunology, viral hepatitis, viral clearance, intracellular cytokine staining, flow cytometry, epitope identification

## Abstract

**Background and aims:** CD4 T cells shape the neutralizing antibody (nAb) response and facilitate viral clearance in various infections. Knowledge of their phenotype, specificity and dynamics in hepatitis E virus (HEV) infection is limited. HEV is enterically transmitted as a naked virus (nHEV) but acquires a host-derived quasi-envelope (eHEV) when budding from cells. While nHEV is composed of the open-reading-frame (ORF)-2-derived capsid, eHEV particles also contain ORF3-derived proteins. We aimed to longitudinally characterize the HEV-specific CD4 T cells and neutralizing antibodies that target either nHEV or eHEV particles in immunocompetent individuals with acute and resolved HEV infection.

**Methods:** HEV-specific CD4 T cells were analyzed by intracellular cytokine staining after stimulation with *in silico* predicted ORF1- and ORF2-derived epitopes and overlapping peptides spanning the ORF3 region. *Ex vivo* multi-parametric characterization of capsid-specific CD4 T cells was performed using customized MHC class II tetramers. Total and neutralizing antibodies targeting nHEV or eHEV particles were determined.

**Results:** HEV-specific CD4 T cell frequencies and antibody titers are highest in individuals with acute infection and decline in a time-dependent process with an antigen hierarchy. HEV-specific CD4 T cells primarily target the ORF2-derived capsid, which correlates with the presence of nAbs targeting nHEV. In contrast, ORF3-specific CD4 T cells are hardly detectable and eHEV is less efficiently neutralized. Capsid-specific CD4 T cells undergo memory formation and stepwise contraction, accompanied by dynamic phenotypical and transcriptional changes over time.

**Conclusion:** The viral capsid is the main target of HEV-specific CD4 T cells and antibodies in acute resolving infection, correlating with efficient neutralization of nHEV. Capsid-specific immunity rapidly emerges followed by a stepwise contraction for several years after infection.

**Impact and implications:** The interplay of CD4 T cells and neutralizing antibody responses is critical in the host defense against viral infections, yet little is known about their characteristics in hepatitis E virus (HEV) infection. We conducted a longitudinal study of immunocompetent individuals with acute and resolved HEV infection to understand the characteristics of HEV-specific CD4 T cells and neutralizing antibodies targeting different viral proteins and particles. We found that HEV-specific CD4 T cells mainly target the viral capsid, leading to efficient neutralization of the naked virus (nHEV) while the quasi-envelope (eHEV) particles are less susceptible to neutralization. As individuals with pre-existing liver disease and immunocompromised individuals are at risk for fulminant or chronic courses of HEV infection, these individuals might benefit from the development of vaccination strategies which require a detailed knowledge of HEV-specific CD4 T cell and antibody immunity.

## Introduction

Hepatitis E virus (HEV) is transmitted enterically and is a major cause of acute hepatitis. Its 7.2 kb positive-strand RNA genome contains three open reading frames (ORF1-3). ORF1 encodes the domains mediating genome replication, ORF2 the virus capsid protein, and ORF3 a small protein critical for virus secretion. Seroprevalence of HEV ranges between ten and thirty percent in large parts of Europe [1], making it the most abundant hepatitis virus in most European countries. The vast majority of these infections are asymptomatic and caused by infections with HEV-genotype 3 [2] which is most commonly acquired by consumption of meat-products from infected animals [3]. However, HEV infection can cause morbidity and even mortality through extrahepatic manifestations, such as neurological disorders, and chronic infections in immunosuppressed individuals that can progress to advanced liver disease, cirrhosis and its complications [2, 4].

Adaptive immune responses are centrally linked to the successful elimination of HEV infection, although the precise contributions of the different arms of the adaptive immune system remain to be identified [5, 6]. Indeed, while data from a macaque model have suggested that neutralizing antibodies (nAbs) may facilitate viral clearance as CD8 T cell depletion only delayed viral clearance [7], recent data from humans with chronic infection clearly link the appearance of CD8 T cell responses to the clearance of chronic infection [8]. nAbs might not only be required to mediate viral clearance, but also to protect from re-infection [9]. However, it remains unclear how long they persist after acute HEV-infection and whether they only target the ORF2-derived capsid in humans.

HEV exists in two different forms: First, highly infectious naked particles (nHEV) that are susceptible to neutralizing antibodies targeting ORF2. These naked particles are usually not found in the circulation but are shed into the feces [10]. Second, quasi-enveloped particles (eHEV) which are covered in host-derived membranes, circulate in the blood stream and are likely responsible for extrahepatic dissemination [11]. ORF3, which is translated from the same subgenomic RNA as ORF2 mediates the interaction with the endosomal sorting complexes required for transport (ESCRT)-machinery and is critical for HEV progeny budding into multivesicular bodies [12]. It is unclear to what extend this protein is also targeted by the humoral and cellular adaptive immune system, although it constitutes the only potential viral target on the surface of quasi-enveloped viral particles [13].

The development of nAb-responses is a tightly regulated process that occurs in the germinal centers of lymphoid tissues and requires the presence of highly specialized CD4 T follicular helper (Tfh) cells [14]. These Tfh cells develop early after antigen-specific priming of naive CD4 T cells and subsequently migrate to the germinal centers and circulate in the periphery [15, 16]. Tfh cells secrete IL-21, which provides maturation signals to B cells and are characterized by the expression of CXCR5 and PD-1 on their surface [14]. Currently, very little information is available about the antigen-hierarchy, kinetics, phenotype and maintenance of HEV-specific CD4 T cell responses and their correlation with total HEV-specific IgGs and neutralizing antibodies targeting naked and quasi-enveloped viral particles. In this study, we set out to address these important questions in acutely and resolved HEV infected individuals.

## Results

### MHC class II epitope identification across the three HEV-ORFs

In order to analyze HEV-specific CD4 T cell responses in a cohort of individuals with acute or resolved HEV infection with diverse MHC class II backgrounds, we performed *in silico* prediction analyses of MHC class II epitopes with the different alleles present in our cohort. As outlined in the methods section, we identified a total of 26 potential epitopes, 15 in the ORF1 region, 11 in the ORF2 region but only one epitope in the ORF3 region. The calculated likelihood of the selected individual peptides to bind the different MHC class II alleles is depicted in Supplemental Figure 2. We aimed to comprehensively identify CD4 T cell immunity to both nHEV and eHEV but could only identify one epitope within the protein encoded by ORF3 using *in silico* prediction. Thus, we used overlapping peptides spanning the entire ORF3-protein. We cultured PBMCs from individuals with acute or resolved infection (Table 1) in the presence of pooled peptides and performed re-stimulation prior to intracellular cytokine staining with individual peptides. As shown in the pie charts in Figure 1A, the percentages of individuals who mounted a CD4 T cell response to any of the tested epitopes within each ORF were comparable. However, the percentages of individuals who mounted CD4 T cell responses to ≥ 4 of the tested epitopes were 0 %, 27 % and 10 % for ORF1-, ORF2- and ORF3-derived epitopes, respectively. Taken together, we functionally validated 20 *in silico*-predicted CD4 T cell epitopes with the majority of these responses targeting the ORF2- but also the ORF1 and ORF3 region (Figure 1A).

**Fig. 1.**
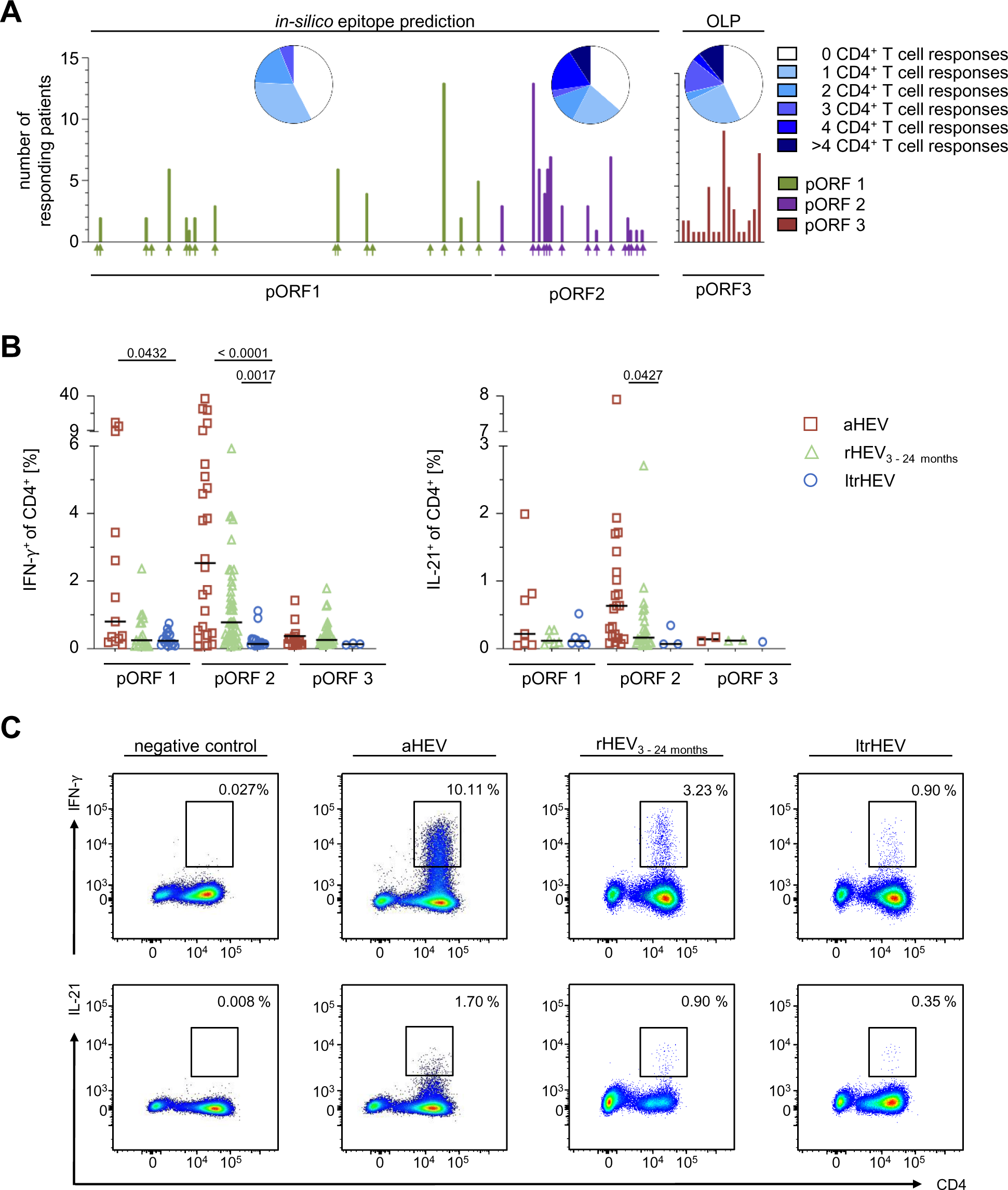
HEV-specific CD4 T cell responses are primarily directed against epitopes of the viral ORF2 protein. PBMCs from individuals with acute or resolved HEV infection were analyzed by ICCS for HEV-specific CD4 T cell secretion of IFN-γ and IL-21 after *in-vitro* culture with HEV peptides identified by *in silico* epitope prediction (ORF1 and ORF2) or overlapping peptides (OLPs) covering the entire ORF3. **(A)** Pie charts display the percentage of individuals with detectable cytokine producing CD4 T cells targeting between 0 and ≥ 4 peptides within each ORF. Below, the number of individuals with detectable cytokine producing CD4 T cells targeting the different epitopes is shown. Colored arrows indicate the localization of the *in silico* predicted epitopes within the ORF1- and ORF2- derived proteins. **(B)** Displayed are frequencies of IFN-γ and IL-21 producing CD4 T cells stratified according to time of infection (acute HEV = red square, resolved HEV_3-24 months_ = green triangle and long-term resolved HEV = blue circle). Levels of significance were calculated by Kruskal-Wallis-Test with *p<0.05% **(C)** Representative pseudocolor dot plots of IFN-γ and IL-21 stainings. Frequencies of cytokine^+^ CD4 T cells are calculated as described previously.

**Table 1:**
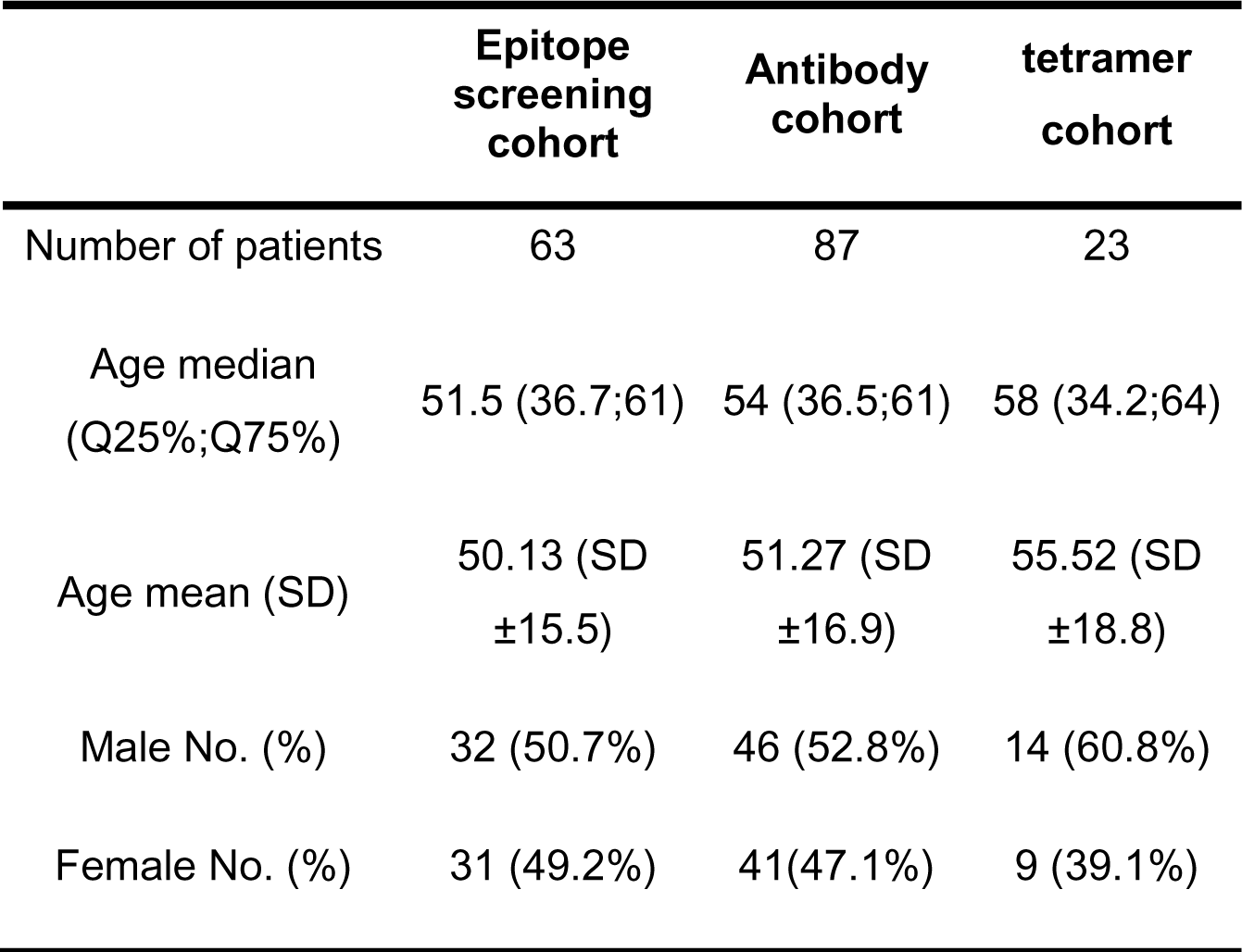
Participant characteristics.

|  | **Epitope**<br>**screening**<br>**cohort** | **Antibody**<br>**cohort** | **tetramer**<br>**cohort** |
| --- | --- | --- | --- |
| Number of patients | 63 | 87 | 23 |
| Age median<br>(Q25%;Q75%) | 51.5 (36.7;61) | 54 (36.5;61) | 58 (34.2;64) |
| Age mean (SD) | 50.13 (SD<br>±15.5) | 51.27 (SD<br>±16.9) | 55.52 (SD<br>±18.8) |
| Male No. (%) | 32 (50.7%) | 46 (52.8%) | 14 (60.8%) |
| Female No. (%) | 31 (49.2%) | 41(47.1%) | 9 (39.1%) |

### Cross sectional analyses of HEV-specific CD4 T cell immunity reveals immunodominance to ORF2

Next, we aimed to analyze the magnitude of the CD4 T cell responses in different phases of acute resolving HEV infection (Figure 1B). We separated our cohort into three groups: Individuals with acute HEV-infection (IgM+) (red), individuals with recently resolved infection within the last 24 months (green) and HEV-seropositive individuals with either unknown time of infection or infection at least 2 years prior to the analysis (blue). We focused our functional CD4 T cell readout (Supplemental Figure 1) on the secretion of IFN-γ and IL-21, the signature cytokines for Th1 and Tfh-biased T cells as well as TNF and IL-2. We observed the strongest responses in terms of IFN-γ and IL-21 production in individuals with acute infection targeting the viral capsid (ORF2), followed by ORF1 and ORF3. Responses against ORF3 were weak, even in the context of acute infection (Fig. 1, B + C). Integrating the percentage of individuals responding to each epitope with the intensity of each individual’s cytokine response, the resulting bubble plots in Figure 2 further visualize the dominance of ORF2-specific CD4 T cells over those targeting ORF1 and ORF3. Importantly, Tfh-biased IL-21 producing CD4 T cell responses were almost exclusively detected within the ORF2-specific CD4 T cells, suggesting strong ORF2-specific B cell helper capabilities of HEV-specific CD4 T cells.

**Fig. 2.**
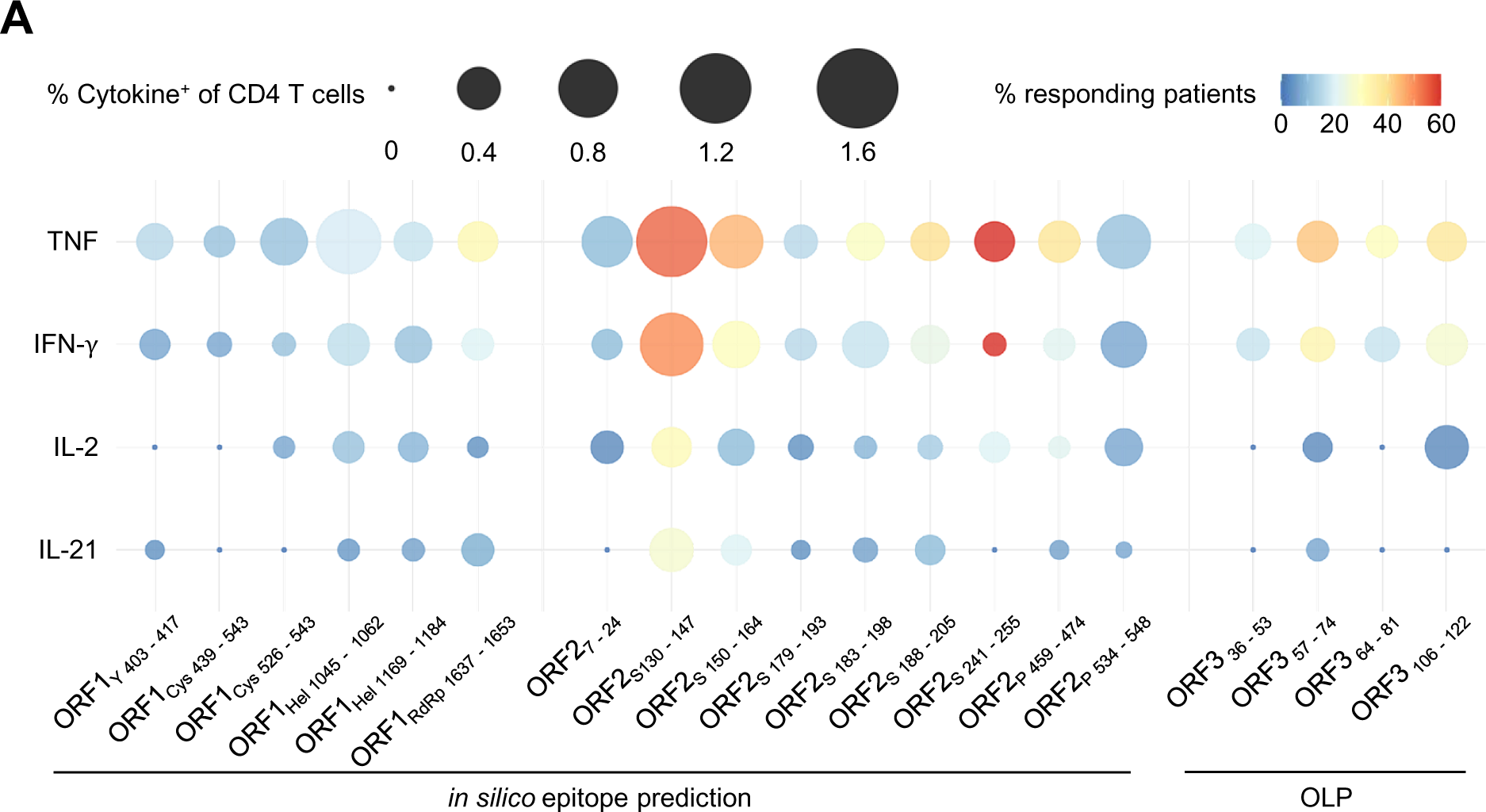
HEV-specific CD4 T cell responses against viral ORF2 are multifunctional. PBMCs from individuals with acute or resolved HEV infection were analyzed by ICCS for HEV- specific CD4 T cell secretion of TNF, IFN-γ, IL-2 and IL-21 after *in-vitro* culture with HEV peptides identified by *in silico* epitope prediction (ORF1 and ORF2) or overlapping peptides (OLPs) covering the entire ORF3. Displayed is a bubble plot illustrating the average magnitude of the HEV-specific CD4 T cell cytokine response (represented by size of the bubbles) and the percentage of individuals mounting a cytokine response (color code) against the most frequently targeted epitopes.

### Cross sectional and longitudinal analyses of HEV-specific humoral immunity against nHEV and eHEV reveals differential neutralization capacity

CD4 T cell help is crucially required to facilitate B cell maturation and antibody secretion. In order to analyze how the HEV-specific CD4 T cell response correlated to the antibody response, we quantified binding and nAbs against HEV. We observed high levels of HEV-specific capsid-binding IgGs in individuals with an infection within the previous five years. In those with an unknown time of infection, anti-HEV IgG titers were significantly lower (Fig. 3A). Anti-HEV IgG titers correlated with the percentage of IL-21 producing ORF2-specific CD4 T cells (Fig. 3B). Nevertheless, the overall correlation was modest, largely owing to the number of individuals with undetectable IL-21 producing CD4 T cells in the group with an unknown time of infection and presumably related to the time of analysis following acute infection. To analyze the neutralization capacities against nHEV and eHEV, we used cell-culture grown naked and quasi-enveloped HEV particles and incubated them with the plasma samples prior to infection and titration on hepatoma cells. Since samples from individuals with acute-viremic infection contain differential amounts of potentially infectious particles skewing our analysis, we only included samples from individuals who had cleared infection. Analysis of nAbs targeting nHEV revealed a strong neutralization capacity early after acute infection followed by a reduction in individuals with an infection time more than three months ago. Neutralization titers then remained stable with a further reduction in individuals with an unknown time of infection (Fig. 3C). This trend was clearly visible in the cross-sectional as well as the longitudinal analyses in individuals with consecutive available plasma samples (Fig. 3D). Again, this was reflected in a correlation of IL-21 producing HEV-specific CD4 T cell responses and the neutralization titers (Fig. 3E). Anti-HEV binding IgG titers also correlated with titers of neutralizing antibodies against naked virions (Fig. 3F). In contrast to the potent neutralization of nHEV using a 1:10.000 serum dilution, analyses of neutralization titers against eHEV with the same dilution factor revealed poor neutralization at all post infection timepoints (Fig. 3G). Taken together, these data demonstrated that ORF2-specific total IgGs and nAbs are present early after acute infection and that they are sustained over several years. In contrast, nAbs against eHEV were hardly detectable during any time after HEV infection.

**Fig. 3.**
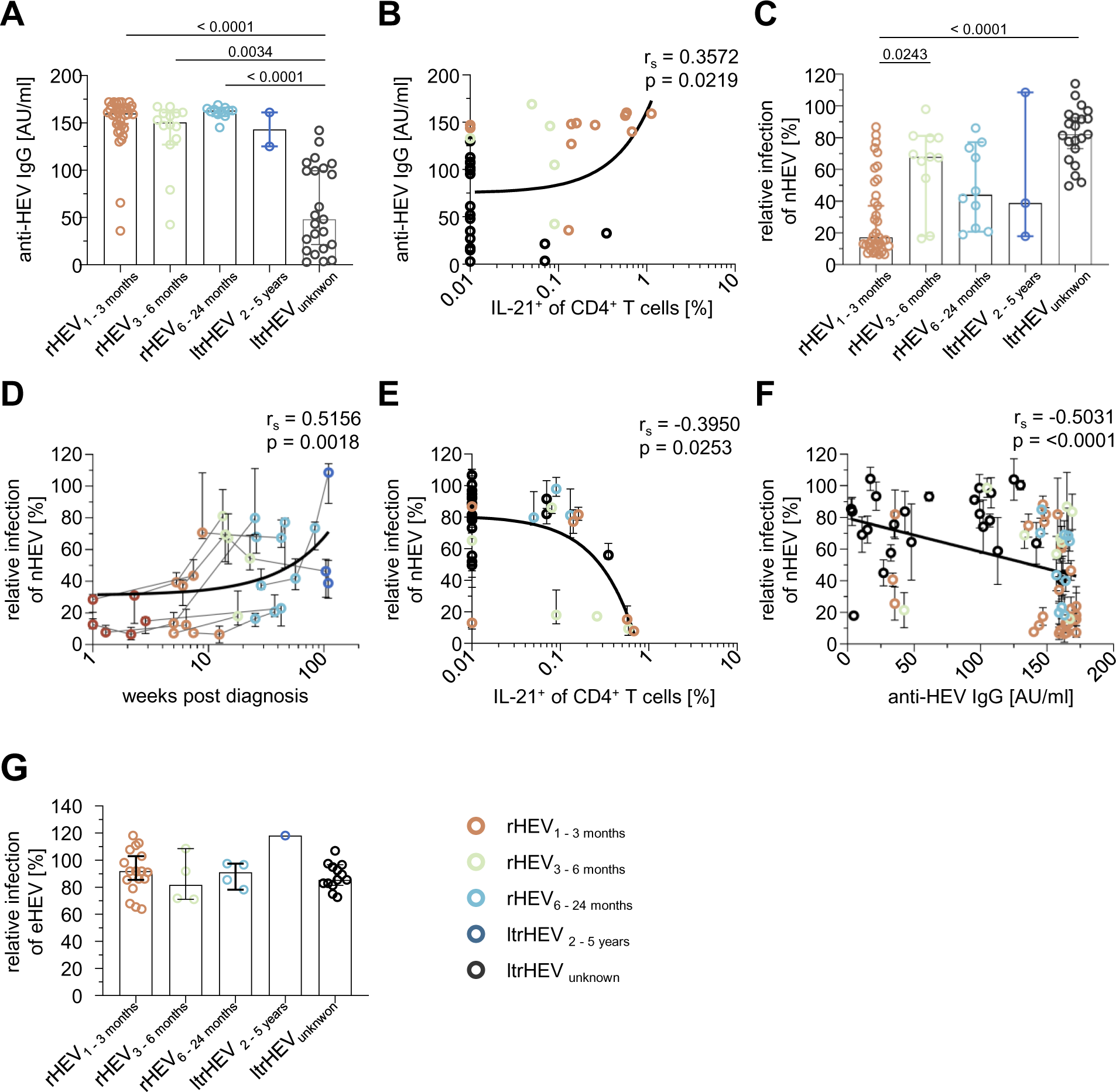
HEV-specific IgG and nAbs against nHEV are maintained for years after viral clearance and correlate with HEV-specific CD4 T cell responses. **(A)** HEV-specific IgG levels in AU/ml (capsid-based ELISA). **(B)** Correlation between HEV-specific IgG levels and IL-21 producing ORF2-specific CD4 T cells. **(C)** The neutralizing capacity against nHEV was determined by analyzing the relative infectivity of target cells with a defined amount of nHEV following co-culture with plasma samples, normalized to controls. **(D**) Longitudinal analysis of HEV neutralization from individuals with 2 or more longitudinal plasma samples. (**E)** Correlation of nHEV neutralization capacity and IL-21 producing ORF2-specific CD4 T cells. **(F)** Correlation of nHEV neutralization capacity and HEV-specific IgG levels. **(G)** The neutralizing capacity against eHEV was determined as described in (C). Statistical significance was determined either by Kruskal-Wallis-Test (A, C, G) or Spearman correlation (B, D-F) and are displayed within figures with *p<0.05. Linear regression was applied. Spearman r (r_s_)

**Table 2:** Clinical data of patients with acute HEV infection.

|  |  |  |
| --- | --- | --- |
| GPT median (MIN;MAX) [U/l] | 197(18;4738) | *14/14 |
| Bilirubin median (MIN;MAX)<br>[mg/dl] | 1.5 (0.2;13) | *14/14 |
| HEV-RNA | Positive | *14/14 |
| HEV-RNA median (MIN;MAX)<br>[IU/ml] | 3500 (429;6100000) | Quantification<br>available in 8/14 |

### Rapid induction and stepwise contraction of HEV ORF2-specific CD4 T cell responses

Next, to better define the role of ORF2-specific CD4 T cell responses and their correlation with ORF2-specific total IgGs and nAbs, we performed an *ex vivo* quantification and phenotypical analysis. In HLA-restriction analyses, we were able to demonstrate that the immunodominant epitope ORF2s130-147 is restricted by HLA DRB1*04:01 and the epitope ORF2s188-205 by HLA-DRB1*01:01 (Fig. 4A). We obtained customized MHC class II tetramers for both epitopes and tested our cohort for *ex vivo* responses against these epitopes (Supplemental Figure 4). Among all individuals within our cohort, 7/11 DRB1*04:01-positive individuals showed a response targeting ORF2s130-147 and 10/12 DRB1*01:01-positive individuals had an ORF2s188-205-specific CD4 T cell response. As expected, the highest *ex vivo* frequencies of HEV-specific CD4 T cells were observed in individuals with acute, viremic infection (Fig. 4, B+C). Interestingly, while the frequency decreased rapidly after acute infection and remained stable for at least 24 months after infection, we observed a further reduction in individuals with unknown time of infection (Fig. 4C). These data demonstrate that HEV-specific CD4 T cell frequencies contract immediately after viral clearance and suggest a second phase of contraction more than two years later.

**Fig. 4.**
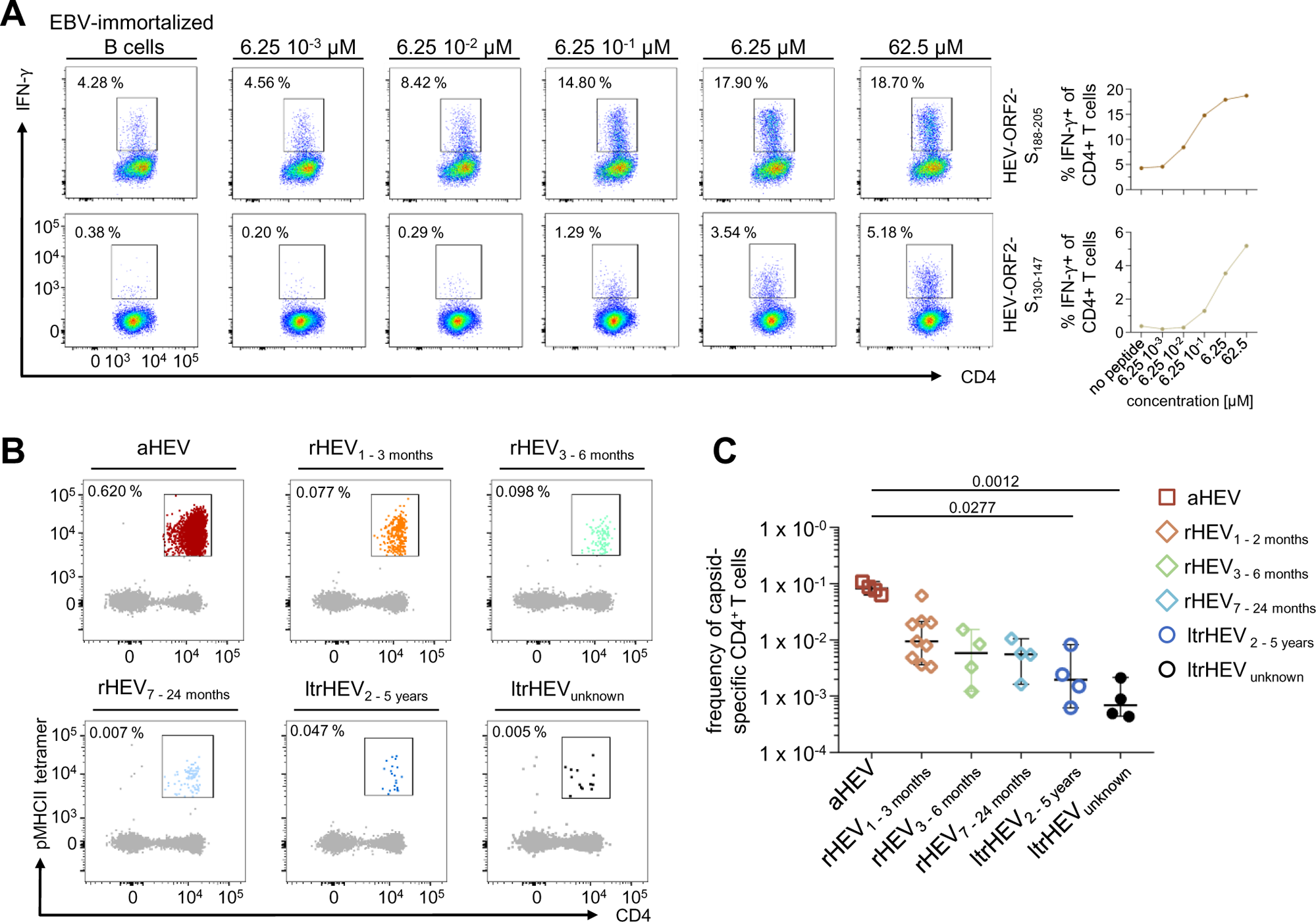
Stepwise contraction of ORF2-specific CD4 T cells following HEV infection. **(A)** Two immunodominant CD4 T cell epitopes were identified by *in vitro* ICCS analysis, both originating from the S-domain of ORF2. HLA-DRB1*-affinity was approximately determined by *in vitro* ICCS experiments using increasing peptide concentrations of HEV-ORF2-S_188-205_ towards HLA-DRB1*01:01 and HEV-ORF2-S_130-147_ towards HLA-DRB1*04:01 respectively. HLA restriction assays were performed using Epstein-Barr-Virus immortalized B cells with carrying the respective HLA class II alleles. **(B)** pMHCII-tetramer enrichment was performed on PBMCs from 23 individuals with acute HEV infection or a confirmed HEV resolved status. (37 samples from 23 individuals were analyzed, tetramer-binding was observed in 29 samples from 17 individuals). Pseudocolor plots showing DRB1*01:01/-S_188-205_ and DRB1*04:01/-S_130-147_-specific CD4 T cells *ex vivo* at different timepoints following acute HEV infection. **(C)** Cumulative ex vivo frequency for DRB1*01:01/-S_188-205_ and DRB1*04:01/-S_130-147_ indicated at different timepoints following acute HEV infection. Kruskal-Wallis-Test was applied with *p<0.05.

### Dynamic phenotypical and transcriptional changes in ORF2-specific CD4 T cell memory

Next, we aimed to analyze the dynamics of the development of CD4 T cell memory and the phenotypic and transcriptional characteristics of the different memory stages in this acute-resolving infection. Dimension reduction of a large flow cytometric dataset including phenotypic markers usually associated with activation (CD38, ICOS, KI-67, PD-1), memory (CD127), transcription factors (T-bet, TOX, TCF-1) and lineage commitment-defining chemokine receptors (CXCR5, CXCR3) revealed a distinct clustering of the ORF2-specific CD4 T cells obtained at different infection timepoints (Fig. 5A). As expected, markers of activation were largely restricted to the acute infection timepoint (Fig. 5, B – D), although PD-1 expression was maintained at high levels over several years after infection and was found to be lower in individuals with unknown time of infection (Fig. 5E). The rapid decline of classical T cell activation markers CD38 and ICOS early after acute infection coincided with an upregulation of the memory markers CD127 and TCF-1 (Fig. 5, F + I) and a loss of expression of the transcription factors T-bet and TOX (Fig. 5, J + K). Expression of the chemokine receptors CXCR5 and CXCR3 can be helpful to analyze T helper lineage commitment (Fig. 5, G + H). While CXCR3 expression is associated with Th1 commitment, Tfh cells typically express CXCR5. Importantly, however, in the memory phase, CXCR5 expression is not restricted to the Tfh compartment and CXCR5-expressing cells can serve as precursor for diverse effector CD4 T cells in a secondary response [17, 18]. We found that both chemokine receptors were more strongly expressed on HEV-specific CD4 T cells from individuals with unknown time of infection compared to those from individuals with documented infection within the last two years. In addition, diffusion map embedding revealed that HEV-specific CD4 T cells from individuals with unknown time of infection localized at the opposing end of the map compared to those from individuals with acute infection (Fig. 6). Collectively, these data suggest that maintenance of HEV-specific memory CD4 T cells is a dynamic process that is associated with phenotypic and transcriptional changes even years after spontaneous viral clearance.

**Fig. 5.**
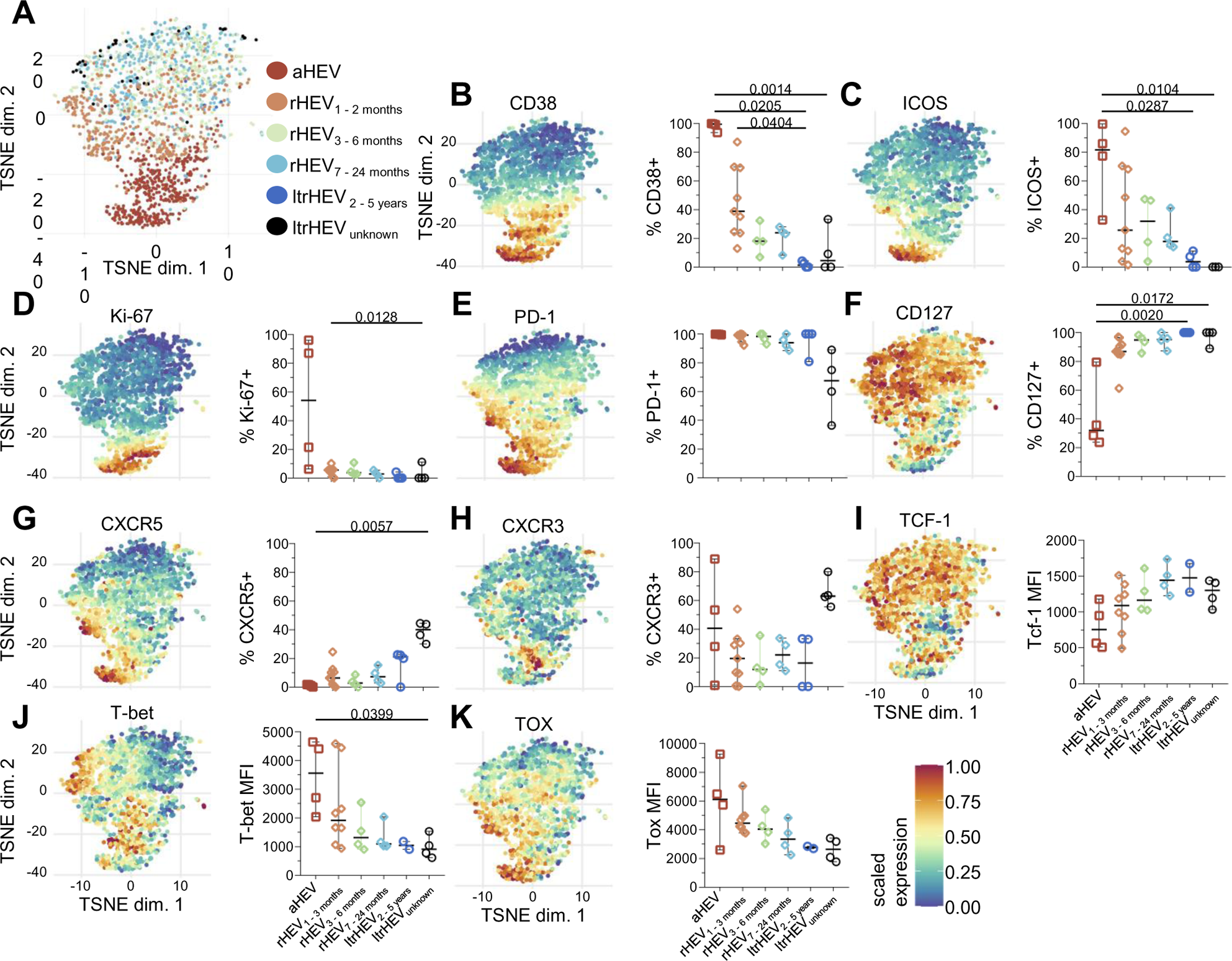
Dynamic phenotypical and transcriptional changes in ORF2-specific CD4 T cell memory. **(A)** Nonlinear dimensionality reduction t-SNE technique was performed on tetramer-binding HEV-specific CD4 T cells from all samples shown in Figure 4C. **(B - H)** Expression levels of the indicated markers on HEV-specific non-naive CD4 T cells at the indicated time post infection. **(I-K)** MFIs of transcriptional factors on HEV-specific non-naive CD4 T cells at the indicated time post infection. Statistical significance was determined by Kruskal-Wallis-Test (B-K) and is displayed within figures with *p<0.05.

**Fig. 6.**
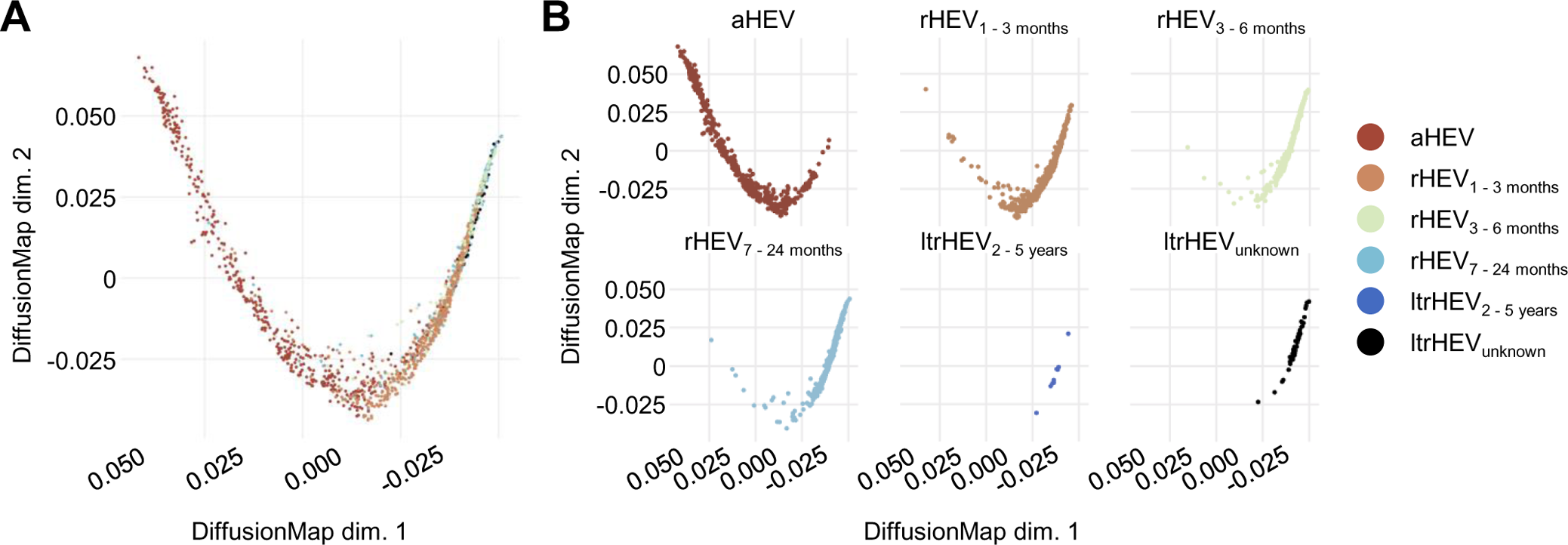
Diffusion mapping confirms continuous memory differentiation of ORF2-specific CD4 T cells. **(A + B)** Diffusion maps used for dimensional reduction of flow cytometric data of DRB1*01:01/_-S188-205_- and DRB1*04:01/-_S130-147_-specific CD4 T cells stained with the markers presented in figure 5. Each dot represents one cell. Aggregate of all samples is displayed in **(A)** and groups stratified by time post infection are displayed in **(B).**

## Discussion

HEV infection is cleared without causing symptomatic disease in most immunocompetent individuals. However, it can induce a variety of clinical events ranging from extrahepatic manifestations in healthy individuals over acute hepatitis or hepatic decompensation in patients with chronic liver disease to persistent infection resulting in liver fibrosis or cirrhosis in immunosuppressed individuals. Despite seroprevalence rates up to 32 % in western countries and its relevance in causing morbidity in vulnerable individuals, little is known about the precise characteristics of HEV-specific CD4 T cell immunity. Our study aimed to provide insights into three central areas of HEV-specific immunity: (1) The immunological targets of naked and quasi-enveloped virions and the associated proteins, (2) the correlation between HEV-specific CD4 T cell immunity and antibody responses and (3) the longevity of HEV-specific antibodies and CD4 T cells and their memory characteristics over time after successful viral clearance.

Previous studies have shown that the viral capsid, encoded by ORF2, is an important target of the antiviral CD4 T cell [19] and antibody response [20], although it has not unanimously been identified as the dominant target of T cell immunity [21]. In HEV-infected macaques, the occurrence of ORF2-specific antibodies is closely linked to viral clearance suggesting that ORF2-specific antibodies are centrally involved in mediating immune control [7, 9]. It is unclear, however, how this control is facilitated as ORF2-specific antibodies cannot bind serum-derived eHEV [22]. Neutralization within endolysosomes of the target cells after degradation of the quasi-envelope has been suggested as a mechanism [23]. It remains unclear, however, whether quasi-enveloped virions are completely shielded from immune attacks. Indeed, in vitro studies have shown that eHEV can be neutralized by ORF3-specific antibodies, although this process is not very efficient [13, 23]. In the macaque model, eHEV neutralization has also been described [7]. As the quasi-envelope is derived from a host membrane built around the viral protein encoded by ORF3, this protein is the only virus-derived structure that is potentially accessible to neutralizing antibodies [23]. We found that CD4 T cells targeting ORF3-derived epitopes were significantly lower in frequency compared to those targeting ORF2-specific epitopes and that they were barely detectable even in acutely infected individuals. The most likely explanation for this might be that the ORF3-derived protein does not include amino acid-sequences that are readily presented by HLA class II molecules as we could not identify potentially strong binding ORF3-derived epitopes *in silico*. We also could not detect IL-21 production in ORF-3-specific CD4 T cells, a Tfh cell-derived cytokine that facilitates B cell maturation and generation of antibody-producing plasma cells. The lack of a strong ORF3-specific CD4 T cell response, particularly the absence of IL-21 production, might partly explain the poor neutralization of eHEV that was observed in our study. Indeed, although some degree of neutralization could be observed with high serum concentrations (Supplemental Fig. 3), nHEV was readily neutralized at a 1:10.000 serum-dilution while eHEV neutralization could not be observed with this serum dilution. The inability of the immune system to mount responses against eHEV might also contribute to the virus’ ability to establish persistent infection in immunosuppressed hosts. In contrast, we observed high frequencies of ORF-2-specific CD4 T cells in acutely infected individuals that declined over time but were maintained over years and were able to produce IL-21. This observation correlated with the presence and maintenance of ORF-2 specific binding and neutralizing antibodies. Thus, characteristics of CD4 T cells and antibodies targeting the different forms of the virus were closely linked and we could demonstrate that HEV-specific immunity is dominated by ORF-2-specific immune responses.

Although ORF-2-specific antibodies cannot directly target or opsonize eHEV that circulates in the serum, they are central for prevention from productive HEV re-infection [9]. It is unclear however, how long sterilizing immunity persists as cases of re-infection have been described in seropositive individuals [24, 25]. We therefore aimed to study the longevity of adaptive HEV-specific immune responses and were able to demonstrate that antibody levels and HEV-specific CD4 T cell frequencies remain high for several years after infection. However, in individuals with unknown time of infection, both CD4 and antibody levels were significantly lower, suggesting that HEV-specific adaptive immunity is not maintained at stable levels over decades, as has been shown for other antiviral immune responses such as smallpox [26, 27]. Using customized tetramers bearing newly identified MHC class II epitopes, our study is the first to longitudinally analyze *ex vivo* frequencies of HEV-specific CD4 T cells. Our data suggest two phases of contraction within the HEV-specific CD4 T cell compartment. After the first contraction phase following viral clearance, we observed rather stable frequencies of HEV-specific CD4 T cells in individuals with a documented time of infection within the last 2 to 3 years. Individuals with an unknown time of infection had even lower frequencies, suggesting a stepwise contraction several years after viral clearance. We did not only observe a reduction in HEV-specific CD4 T cell frequencies years after viral clearance, but this reduction was also associated with phenotypic changes within HEV-specific CD4 T cells. While well-established markers of T cell memory, such as CD127 and TCF-1 [17] were strongly expressed at all late time points after acute infection, suggesting a formation of stable memory, we observed phenotypic changes that included a downregulation of PD-1 and higher expression of the chemokine receptors CXCR5 and CXCR3 in individuals with an unknown time of infection, suggesting that HEV-specific memory CD4 T cells retain developmental plasticity. It will be important to determine in future studies which of the memory T cell clones are maintained in the second contraction phase and what the functional implications of the changes in the memory compartment are.

The relative contribution of the different arms of the immune system to viral control in HEV infection remain an area of debate. While CD8 T cells have been found to be major players in facilitating immune control to HBV and HCV infections [28, 29], depletion of CD8 T cells only delayed viral control by a week in HEV-infected macaques [7]. Instead, viral clearance in CD8-depleted macaques was associated with a robust CD4 T cell and antibody response, suggesting a central role for antibodies and CD4 T cells in viral clearance. The strong CD4 T cell responses and the high titers of neutralizing antibodies in acutely infected individuals in our study support this notion and provide clear evidence for the first time that HEV-infection induces CD4 T cells that are able to provide B cell help by producing IL-21 in the acute phase of the infection and subsequently undergo memory differentiation that might protect from secondary infections. The observation that CD8 T cells might be dispensable, however, could be specific to resolution of acute HEV-infection. Indeed, the appearance of HEV-specific CD8 T cells in individuals chronically infected with HEV is linked to immune control and resolution of persistent infection [8].

The central limitation of our study is the lack of precise information on the time of infection in seropositive individuals with unknown time of infection. While the waning antibody titers, the reduced frequency of HEV-specific CD4 T cells and the diffusion mapping of the integrated phenotyping of tetramer-positive cells suggest that the time of infection is longer ago than in the cohort with a known time of infection, we cannot state with absolute certainty that infection is indeed longer ago than five years.

Collectively, our data show that HEV-specific CD4 T cell and humoral immunity primarily targets the viral ORF2-derived capsid and naked virions. Functional analyses of HEV-specific CD4 T cells demonstrate Tfh-characteristics of ORF2-specific CD4 T cells through IL-21 production and MHC-class II tetramer analysis show memory formation of HEV-specific CD4 T cells while maintaining phenotypic plasticity.

## Methods

### Study cohort

A total of 126 individuals exposed or acutely infected with HEV were screened for inclusion in this study. Individuals were HLA typed by next generation sequencing (NGS) using commercially available primers (GenDx) and run on a MiSeq system. Acquired data was analyzed using the NGSsengine software (GenDx). A total of 88 individuals were included in this study. Plasma samples and PBMCs were frozen until usage. Characteristics are summarized in supplementary table 1.

This study was approved by the Ethics Committee of the Albert-Ludwigs-University Freiburg (474/14, 201/17, 486/19), and informed, written informed consent was obtained from all blood donors before enrollment in the study.

### Synthetic peptides

26 peptides originating from ORF 1, 2 were designed using the IEDB *in silico* epitope prediction tool (https://www.iedb.org/home_v3.php). Additionally, 16 overlapping peptides (OLPs) were designed spanning the whole amino acid sequence of ORF 3 in 18-mers. Due to technical difficulties OLP 10 could not be synthesized as an 18-mer and was therefore ordered as a 14-mer. Additionally six prediscribed CD4 T cell peptides of the Hepatitis E Virus were synthesized. All peptides are listed in supplementary table 3 and 4 and were synthesized by Genaxxon Bioscience with >70% purity.

### Preparation of peripheral blood mononuclear cells for staining or *in-vitro* expansion

Peripheral blood mononuclear cells (PBMCs) were isolated from EDTA-whole blood samples after density gradient centrifugation using Pancoll (Pan Biotech, Germany) and washing with phosphate-buffed saline (D-PBS ThermoFischer, Germany). Subsequently, the isolated cells were cryopreserved in a dimethyl sulfocide (DMSO) freezing solution at −80°C. Once needed the PBMCs were thawed using a complete medium (RPMI1640, 10% fetal calf serum, 100 U/ml penicillin, 100 µg/ml streptomycin, 10 mM HEPES; TermoFischer, Germany).

### *In-vitro* expansion

The *in-vitro* expansion was performed as described previously with 2 x 10^6^ PBMCs [30]. In brief, PBMCs were stimulated with HEV peptides (10µM) for 14 days. Initially an agonistic CD28 mononuclear antibody (αCD28mAb) (10µg/ml) was added to the culture. After 14 days of culture, a single peptide re-stimulation with an ICCS was conducted. Media and additive were changed on days 3, 7 and 10.

### Intracellular cytokine staining (ICCS)

After 14 days of *in-vitro* T cell expansion approximately 2.5 x 105 cells, per well, were re-stimulated with single HEV-peptides (7,5µM), phorbol myristate acetate (PMA) (20ng/ml; Sigma-Aldrich Chemie GmbH, Germany) and ionomycin (1µg/ml; Sigma-Aldrich Chemie GmbH, Germany) (positive control) or left unstimulated (negative control) for 5 hours. Additionally, 0,5 µl/ml Brefeldin A and 0,325µl/ml Monensin (BD Biosciences, Germany) were added to each well. Afterwards cells were stained for 15 minutes at room temperature with surface markers, then fixed and permeabilized (Cytofix/Cytoperm, BD Biosciences, Germany) and stained with antibodies against cytokines for 30 minutes at room temperature. Antibodies are listed in supplementary table 2. Fixable Viability dye^eFluor780^ (eBioscience, Germany) was used to discriminate live from dead cells. Samples were acquired with a CytoFLEX S (Beckman Coulter, Germany) and a FACSCanto (BD Biosciences, Germany) flow cytometer and analyzed with FlowJo X 10.0.7r2 software (LLC, BD Life Sciences, USA). Gating was performed as described previously [30]. For the analysis dead (viability dye^+^) cells were excluded.

### HLA-DRB1 epitope restriction assay

Human leukocyte antigen restriction assay was performed using HLA-DRB1*01:01 or DRB1*04:01-carrying Epstein-Barr-Virus immortalized B cells (EBBs), which pose as antigen presenting cells (APCs) for epitope pre-stimulated CD4 T cells. To exclude non-specific binding of the tested peptides, we used EBBs with different HLA-DRB1 molecules, e.g. DRB1*15:01, DRB1*11:01, DRB1*03:01 as controls. On day 13 approximately 2.5 x 10^6^ EBBs per well, were given single HEV-peptides with different concentration (0.0065 µM, 0.0625 µM, 0.625 µM, 6.25 µM, 62.5 µM and 0 µM peptide concentration respectively) and incubated overnight.

The PBMCs *in-vitro* expansion was performed as described previously for individuals carrying either HLA-DRB1*01:01 or DRB1*04:01. On day 14 EBBs and PBMCs co-cultures matching the HLA-DRB1 genes were created. The cells were then incubated for 5 hours before ICCS was conducted.

### Magnetic bead-based enrichment of antigen-specific CD4 T cells

The magnetic bead-based Tetramer enrichment of antigen-specific CD4 T cells was performed according to previously described protocols [31, 32]. Anti-PE magnetic beads were used according to the manufacturer’s instructions (MACS Technology, Miltenyi Biotec). The customized PE-labeled tetramers were obtained from MBL life sciences (via Biozol, Germany). In brief, PBMCs from HLA-DRB1*01:01 or DRB1*04:01 positive donors were incubated with the respective MHC class II epitope-specific tetramer (supplementary table 4). HEV-specific cells were enriched by magnetic cell separation after adding anti-PE magnetic beads. Both the enriched and the depleted sample, as well as the pre-enriched sample were stained with fluorochrome-conjugated antibodies and analyzed by flow cytometry. All samples were acquired using an LSRFortessa flow cytometer (BD Biosciences, Germany) and analyzed with FlowJo X 10.0.7r2 software (LLC, BD Life Sciences, USA). For the analysis of HEV-specific CD4 T cells doublets were excluded and bulk CD4 T cells were gated on dump negative (CD14^+^, CD19^+^, viability dye^+^) cells. Naïve cells were distinguished by CCR7^+^ and CD45RA^+^. Tetramer^+^ cells were then gated on CD4^+^ non-naïve cells as described previously [33]. Frequencies of HEV-specific CD4 T cells were calculated as follows: Absolute number of Tetramer^+^ CD4 T cells (enriched sample) divided by the absolute number of CD4 T cells (pre-enriched sample) times 100. Samples with less than five antigen-specific CD4 T cells were excluded from the final analysis.

### Surface staining and intranuclear transcription factor staining

Staining of surface markers was performed for 20 minutes at 4°C with antibodies listed in supplementary table 2. Cells were fixed and permeabilized with eBioscience Intracellular Fixation & Permeabilization Buffer Set (Thermo Fischer Scientific, Germany) and stained with intranuclear antibodies against transcriptional factors at 4°C for 30 minutes (see supplementary table 2). All samples were acquired using an LSRFortessa flow cytometer (BD Biosciences, Germany) and analyzed with FlowJo X 10.0.7r2 software (LLC, BD Life Sciences, USA).

### Dimensional reduction of multiparametric flow cytometry data

Dimensionality reduction was conducted with R version 4.1.1 using Bioconductor (release 3.13) CATALYST (Version 1.16.2). The analyses were performed on HEV-specific live non-naïve CD4 T cells including the markers CD38, ICOS, Ki-67, PD-1, CD127, Tcf-1, T-bet, TOX, CXCR5 and CXCR3. Downsampling to 450 cells was performed prior to dimensionality reduction. Marker intensities were transformed by arcsinh (inverse hyperbolic sine) with a cofactor of 150. Dimensionality reduction on the transformed data was achieved by t-SNE.

### Plasmids and cells

Plasmid encoding HEV GT3 Kernow-C1/p6 (GenBank accession No: JQ679013) and human hepatoma S10-3 cells were kindly provided by Suzanne U. Emerson, NIH. S10-3 cells were grown in Dulbecco’s Modified Eagle’s Medium (Gibco, Carlsbad, CA) supplemented with 10% fetal bovine serum (FBS; Merck, Darmstadt, Germany) and 1% penicillin-streptomycin (Gibco).

### Production of eHEV and nHEV particles

HEV GT3 Kernow-C1/p6 RNA was produced in-vitro transcription using the mMESSAGE mMACHINE™ T7 Transcription Kit (Life Technologies). 10 µg of RNA was transfected into 4×10^6 S10-3 cells by electroporation with a Gene Pulser II apparatus (Bio-Rad) in 0.4 cm Gene Pulser cuvettes (Bio-Rad) at a capacity of 0.975 nF and a voltage of 0.27 V. For eHEV production, the culture medium was changed 24 hours before harvesting. On day 7 post-electroporation, the supernatant containing eHEV was collected and filtered through a 0.45 um pore size filter (Whatman GE Healthcare Life Sciences).

For nHEV production, the cells were harvested in PBS 7 days post-electroporation and lysed by repeated freeze-thaw cycles in liquid nitrogen. The crude virus was purified by ultracentrifugation at 32000 rpm for 3 hours, at 4°C on a 20% w/v sucrose cushion. The virus pellet was dissolved in PBS and stored at −80°C.

### Neutralization of nHEV virions

5 x 10^3^ S10-3 cells per well were seeded into 96-well plates. The infection with nHEV was conducted in DMEM supplemented with 10% FBS and 1% penicillin-streptomycin. Patient plasma and serum samples were diluted 10^−2^, 10^−3^, 10^−4^, 10^−5^ and incubated with nHEV at a multiplicity of infection (MOI) of 0,8 at 37°C for 15 minutes, before the solution was added to the cells. 24 hours post-infection, the virus was removed, and culture medium was replenished (Supplementary Figure 3).

7 days post infection, the cells were fixed in 4% paraformaldehyde (EMS, Hatfield, PA) and stained as described previously [34] using an anti-ORF2 monoclonal antibody (1:400, 1E6; Millipore, Burlington, MA) and a secondary anti-mouse antibody conjugated to Alexa Fluor 594 (1:500; Thermo Fisher). HEV infection was analyzed as described before [35] using automated microscopy and image analysis with KNIME.

### Neutralization of eHEV virions

4 x 10^4^ S10-3 cells per well were seeded into 48-well plates. The infection with eHEV was conducted in DMEM supplemented with 10% FBS and 1% penicillin-streptomycin. Patient plasma and serum samples were diluted 10^−2^, 10^−3^, 10^−4^ and incubated with eHEV at a MOI of 1×10^−3^ at 37°C for 60 minutes, before the solution was added to the cells. As a control antibody, a polyclonal rabbit anti-ORF2 antibody (a kind gift from XJ Meng, Virginia Tech) was used in a dilution of 20^−2^. 8 hours post-infection, the virus was removed, and culture medium was replenished. 7 days post infection, the cells were fixed and stained as described above. Relative infection analysis was conducted at the cell discoverer 7 microscope (Zeiss) and by counting foci.

### Statistical analysis

Statistical tests were performed using GraphPad 9 software (GraphPad Prism Software Inc., USA). Due to limited sample size, non-parametric tests were used. For the comparison of up to three unpaired groups, Mann-Whitney U test was applied, whereas for unpaired data multiple comparison analyses were performed using Kruskal-Wallis. In case of multiple comparisons Bonferroni correction adjusted the level of significance. The adjusted levels of significance are displayed in the respective figure legends. Correlation analyses were performed by Spearman test. P values less than 0.05 were considered to be significant with *p<0.05; **p<0.01; ***p<0.001; ****p<0.0001.

## Acknowledgements

This work was supported by grants from the Deutsche Forschungsgemeinschaft (DFG, German Research Foundation): project 272983813 (TP01 to RT, TP02 to CNH, TP04 to TBo, TP20 to MH, TP21 to BB and TP22 to VLDT) and project 256073931 (to CNH). TBr also received funding from the DFG (BR 4182/3-1 and SFB1382 Project ID 403224013/B07). The German Center for Infection Research (DZIF), TTU Hepatitis also supported this work - Project 05.823 to VLDT.

## Conflict of interests

All authors will provide a COI form to exclude conflict of interests.

## Data Availability Statement

Data are available upon reasonable request to the corresponding author.

## Authors’ contributions

Study concept and design: BC, MSM, TB; acquisition of data: BC, MSM, LM, LK, JW, KB, KZ, MP; analysis and interpretation of data: BC, MSM, LM, MH, VLDT, TB; drafting of the manuscript: BC, MSM, TB; critical revision of the manuscript for important intellectual content: PR, TBr, BB, CNH, MH, RT, VLDT; statistical analysis: BC, MSM; obtained funding: TBr, BB, CNH, MH, RT, TB; technical, or material support: PR, TBr, VLDT; study supervision: RT, TB.

## Abbreviations

APCs: antigen presenting cells
DMSO: dimethyl sulfocide
EBBs: Epstein-Barr-Virus immortalized B cells
eHEV: quasi-enveloped HEV
ESCRT: endosomal sorting complexes required for transport
HEV: hepatitis E virus
ICCS: intracellular cytokine staining
mAb: mononuclear antibody
nAb: neutralizing antibody
nAbs: neutralizing antibodies
NGS: next generation sequencing
nHEV: naked HEV
OLPs: overlapping peptides
ORF: open-reading-frame
PBMCs: peripheral blood mononuclear cells
PBS: phosphate-buffed saline
PMA: phorbol myristate acetate
Tfh: T follicular helper cells

## Notes

### Competing Interest Statement

The authors have declared no competing interest.

