## Supplements for "Efficient formation and maintenance of humoral and CD4 T cell immunity targeting the viral capsid in acute-resolving hepatitis E infection"

### Supplementary Figures:

#### Supplementary Figure 1

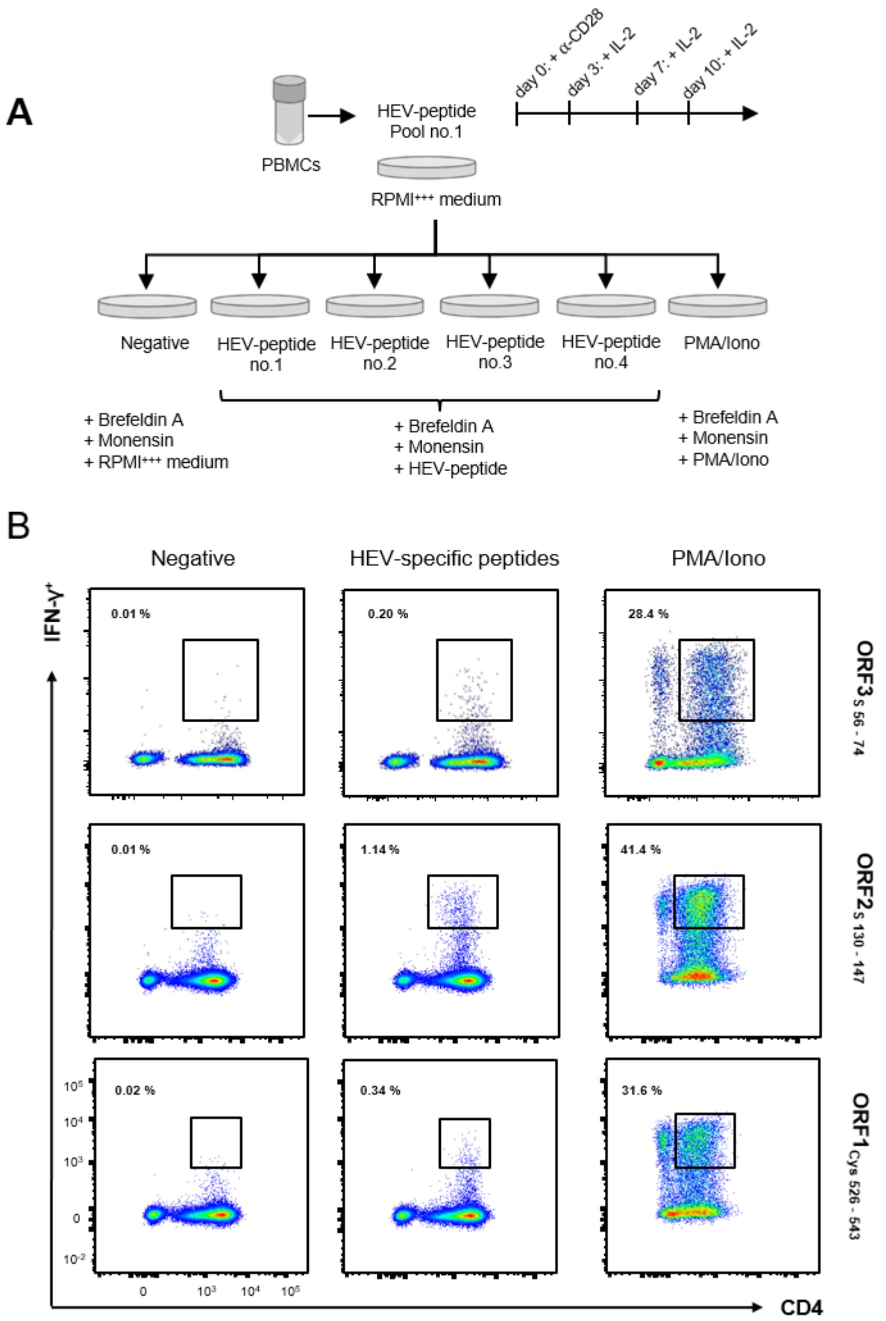


**Supplementary Fig.1 │HEV-peptides are capable of inducing CD4 T cell cytokine response via *in vitro* T cell expansion (A)** Schematic illustation of workflow of *in vitro* expansion and ICCS. HEV-specific CD4 T cell were analysed by ICCS after 14 days of expansion and restimulation with HEV-specific peptides. All HEV-peptides were grouped and different expansion pools of 3 or 4 HEV-specific peptides were generated. On day 14 every culture pool was split and re-stimulation with single peptides followed. HEV-specific CD4 T cell responses were analysed by ICCS for IFN-y and IL-21 production. Negative stimulation and PMA/Iono served as controls. **(B)** Representative pseudo-color plots for negative control (neg), positive control with PMA/Iono simulation and peptide-stimulated originating from each ORF on live single-cell lymphocytes.

Supplementary Figure 2

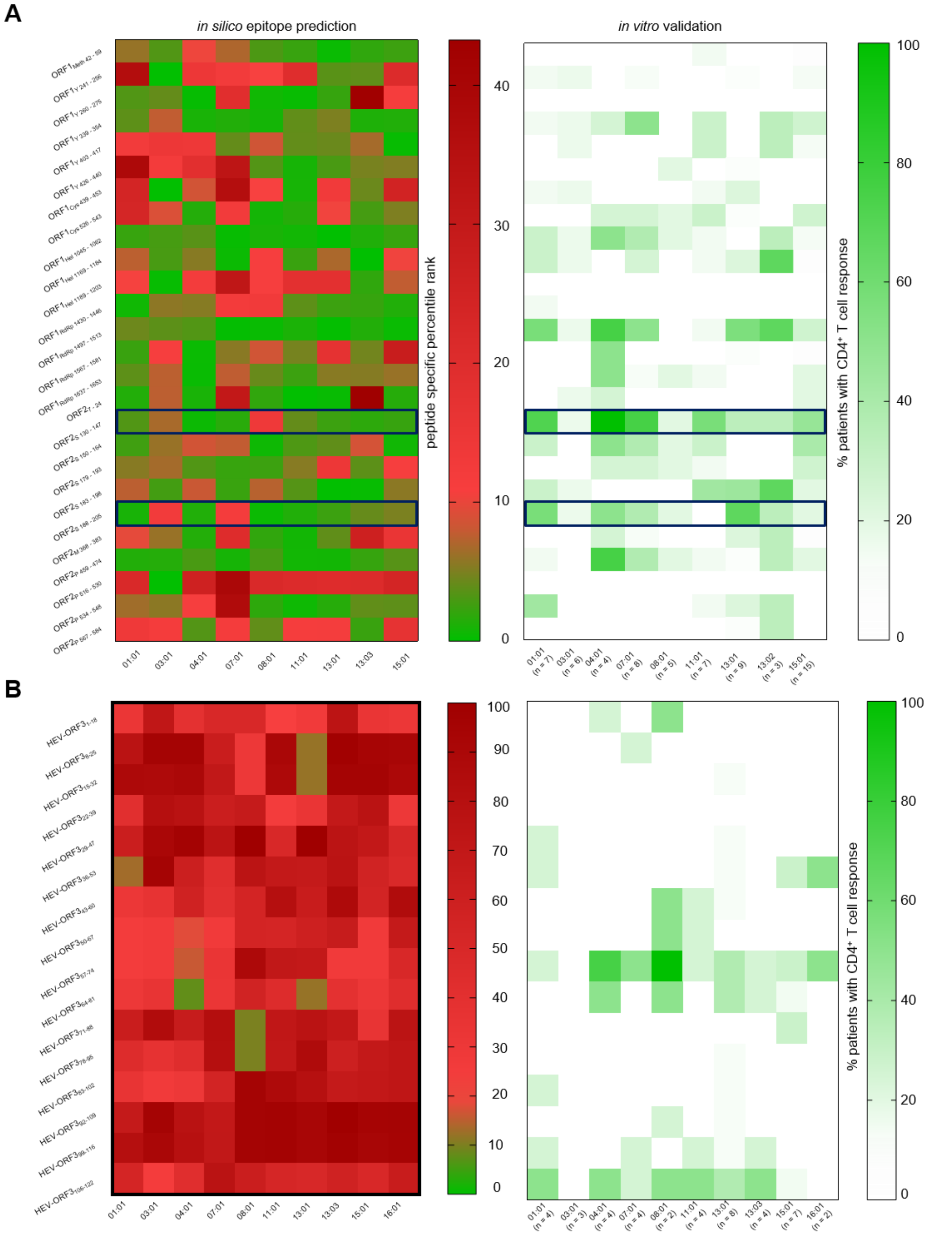


**Supplementary Fig.2 │ *In silico* predicted HLA-DRB1 binding HEV-peptides are recognized by patients carrying different HLA-DRB1 alleles with an acute or resolved HEV infection (A+B)** Heat map of *in silico* prediction for different HLA-DRB1 alleles (left) and *in vitro* validation determined by stimulation of PBMC's with corresponding HEV-specific peptides. The percentage of responding patients for each peptide and each HLA allele is shown (right) **(A)** *In silico* prediciton and *in vitro* validation for 26 HEV-peptides originating from HEV ORF1 and ORF2 (n=37). Blue boxes indicate the peptide sequences that were used for tetramer generation. **(B**) *In silico* prediciton and *in vitro* validation for 16 ORF3 overlapping peptides (OLPs) using PBMC's from resolved HEV infected patients (n=37).

Supplementary Figure 3

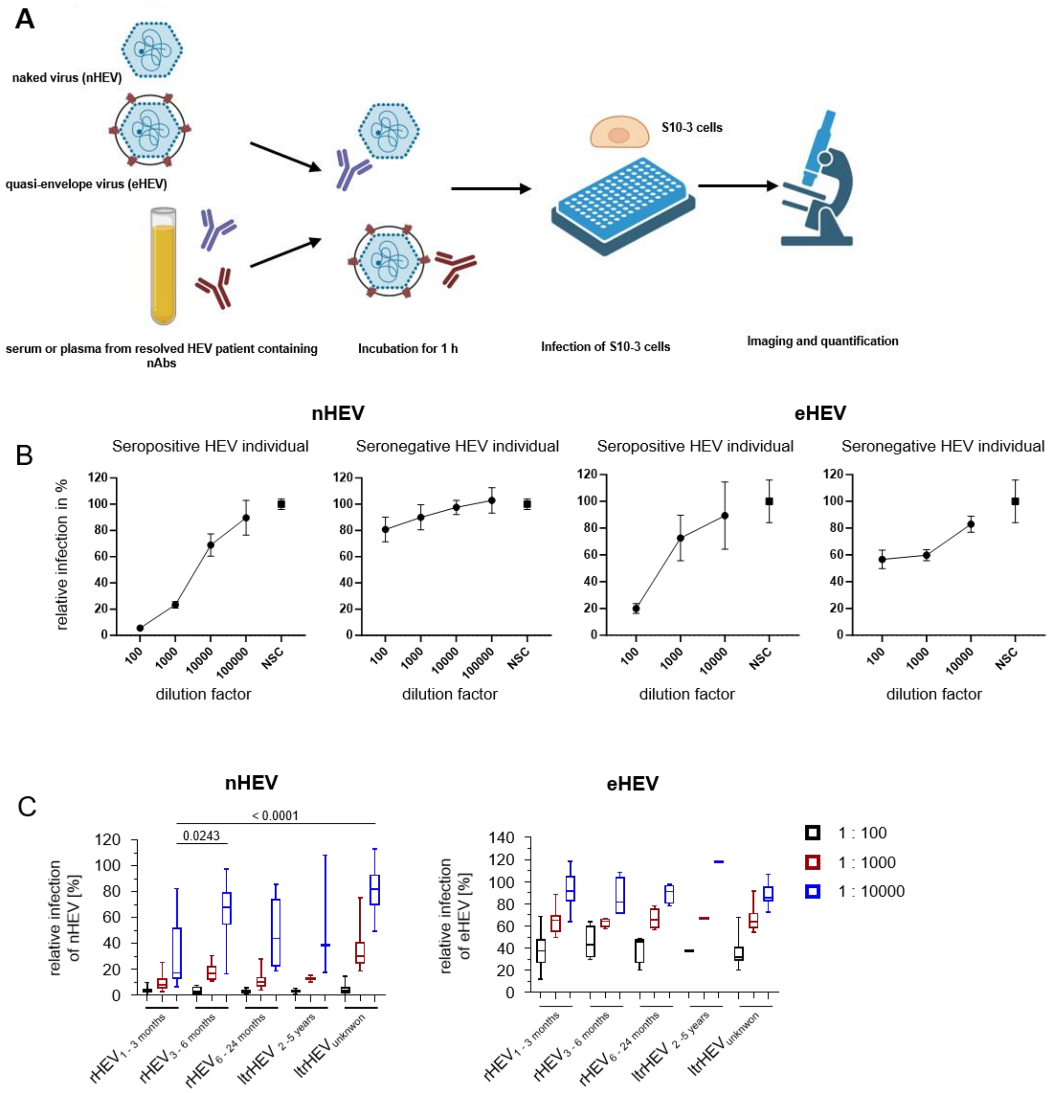


**Supplementary Fig.3 │Neutralization capacity of nHEV is more efficient compared to eHEV. (A)** Serum and plasma samples were collected from 87 resolved HEV patients. The patient's specific neutralizing capacity against either nHEV or eHEV (Kernow-C1 p6 strain) was determined by analyzing HEV infection of S10-3 cells which were incubated with patient serum and nHEV or eHEV using automated microscopy and image analysis. Neutralizing capacity is displayed as relative infection after normalization to controls which were incubated with nHEV or eHEV and without patient serum. **(B)** Neutralization titration representation for nHEV and eHEV. Relative infection against nHEV and eHEV was plotted for different dilution factor of individuals serum (1:100, 1:1000, 1:10.000, 1:100.000 (nHEV) and no-serum control). On the right side. Seronegative patient`s serum served as control. NSC: no-serum control (as normalized to untreated extracellular HEV particle) **(C)** Neutralizing capacity in serum dilutions of nHEV and eHEV at different timepoints after infection. Levels of significance were calculated by Kruskal-Wallis-Test with *p<0.05.

Supplementary Figure 4

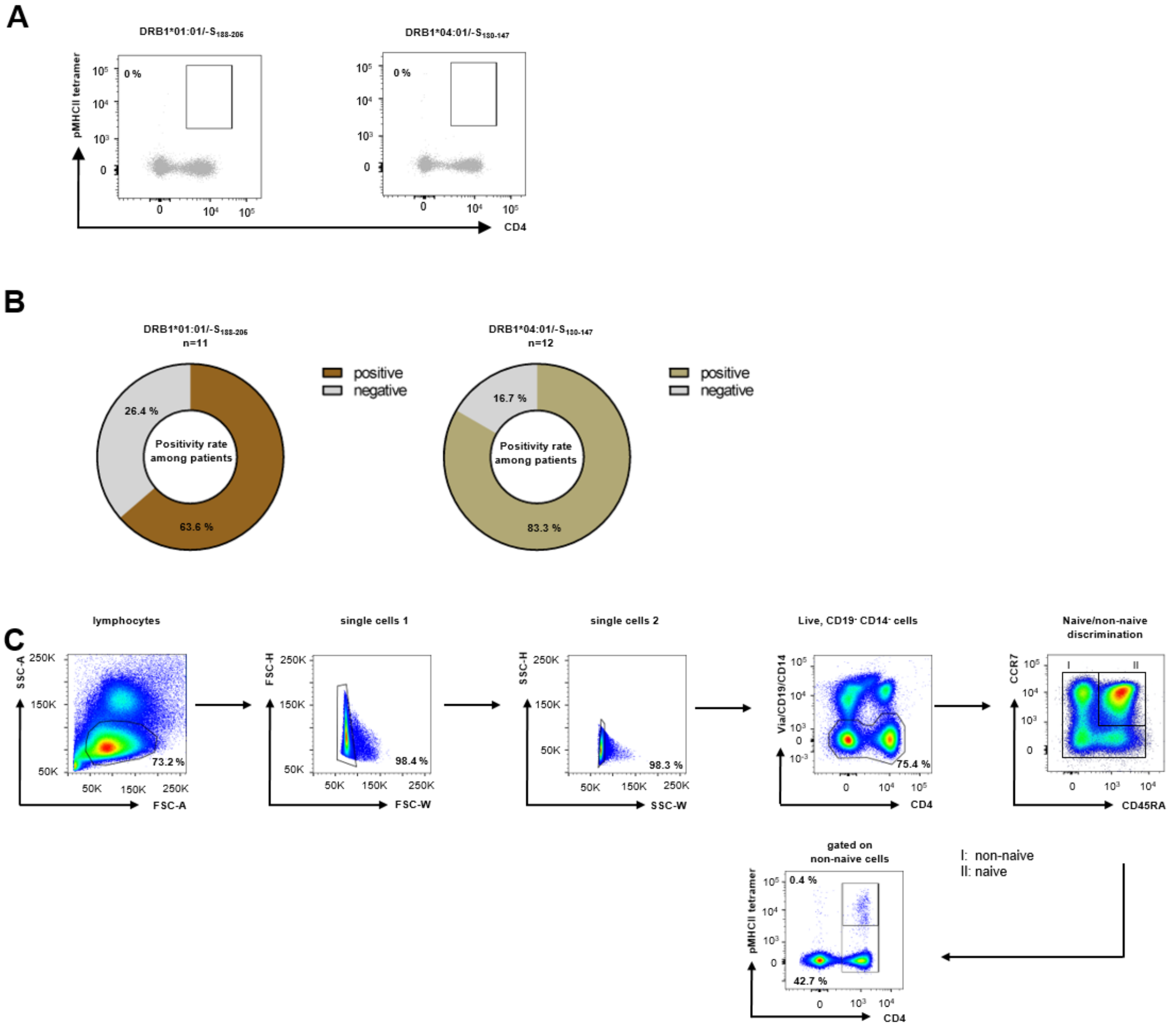


**Supplementary Fig.4 │ORF2-specific CD4 T cells are only detectable in either HLA-DRB1*01:01 or HLA-DRB1*04:01 sero-positive patients (A)** Binding specificity of HEV-specific CD4 T cell pMHCII-tetramers. Both HEV-specific CD4 T cell pMHCII-tetramers were tested each in 3 HEV-seronegative patients by using pMHCII-tetramer enrichment. **(B)** Pie charts representing the number of patients with positive HEV ORF2-specific CD4 T responses for DRB1*01:01/-$S_{188-205}$ (left) and DRB1*04:01/-$S_{130-147}$ (right), respectively. **(C)** Gating strategy of flow cyometry data.

### Supplementary Tables:

Supplementary Table 1
Probe groups

| | Epitope screening Cohort | Antibody Cohort | Tetramer Cohort |
|---|---|---|---|
| aHEV | 11 | 6 | 9 |
| $rHEV_{1-3\ months}$ | 41 | 37 | 9 |
| $rHEV_{3-6\ months}$ | 15 | 17 | 4 |
| $rHEV_{7-24\ months}$ | 10 | 14 | 4 |
| $ltrHEV_{2-5\ years}$ | 4 | 3 | 3 |
| $ltrHEV_{unknown}$ | 40 | 35 | 8 |

Supplementary Table 2
Staining antibodies – Tetramer enrichment and B cell analysis

| Antigen | Fluorophor | Clone | Dilution | Company |
|---|---|---|---|---|
| CCR7 | BUV395 | 3D12 | 1 to 25 | BD Biosciences |
| CD38 | BUV737 | HB7 | 1 to 200 | BD Biosciences |
| CXCR5 | BV421 | RF8B2 | 1 to 100 | BD Biosciences |
| CXCR3 | BV510 | G025H7 | 1 to 33.3 | BioLegend |
| CD127 | BV605 | A019D5 | 1 to 33.3 | BioLegend |
| ICOS | BV711 | DX29 | 1 to 100 | BD Biosciences |
| PD-1 | BV786 | EH12.1 | 1 to 33.3 | BD Biosciences |
| TCF-1 | AlexaFluor488 | C63D9 | 1 to 100 | Cell signaling |
| CD45RA | PerCP-Cy5.5 | HI100 | 1 to 33.3 | Invitrogen |
| T-bet | PE-CF594 | O4-46 | 1 to 33.3 | BD Biosciences |
| Ki67 | PE-Cy7 | Ki-67 | 1 to 200 | BioLegend |
| TOX | APC | TRX10 | 1 to 100 | eBioscience |
| CD4 | Ax700 | RPA-T4 | 1 to 200 | BioLegend |
| Fixable viability dye | eFluor780 | | 1 to 200 | eBioscience |
| CD14 | APC eFluor780 | 61D3 | 1 to 100 | eBioscience |
| CD19 | APC eFluor780 | HIB19 | 1 to 200 | eBioscience |
| CD19 | V500 | HIB19 | 1 to 200 | BD Biosciences |
| IgD | PerCPCy5.5 | IA6-2 | 1 to 50 | BioLegend |
| CD27 | BV605 | L128 | 1 to 100 | BD Biosciences |
| CD20 | PE-Cy7 | 2H7 | 1 to 50 | eBioscience |
| IgG | BV421 | G18-145 | 1 to 50 | BD Biosciences |
| CD3 | eFluor780 | SK7 | 1 to 50 | BD Biosciences |
| IgM | BB515 | G20-127 | 1 to 33.3 | BD Biosciences |
| CD138 | PE | 281-2 | 1 to 20 | BioLegend |
| CD21 | APC | B-ly4 | 1 to 100 | BD Biosciences |
| CXCR5 | BV711 | J252D4 | 1 to 33.3 | BioLegend |
| CD4 | BV510 | SK3 | 1 to 100 | BD Biosciences |
| CD8 | BV421 | RPA-T8 | 1 to 400 | BD Biosciences |
| IFN-g | FITC | 25723.11 | 1 to 8.3 | BD Biosciences |
| IL-2 | PE | MQ1-17H12 | 1 to 25 | BD Biosciences |
| IL-21 | Alexa Fluor 647 | 3A3-N2.1 | 1 to 25 | BD Biosciences |

| Antigen | Fluorophor | Clone | Dilution | Company |
|---|---|---|---|---|
| TNF-a | PE-Cyanine 7 | MAb11 | 1 to 200 | BioLegend |
| Fixable viability dye | eFluor780 | | 1 to 200 | eBioscience |

Supplementary Table 3
HEV Peptides

| Peptide | ORF | Protein | Start | End | Amino acid sequence | Length | Source |
|---|---|---|---|---|---|---|---|
| HEV-ORF1-Meth$_{42-59}$ | ORF 1 | Viral methyltransferase | 42 | 59 | QTEILINLMQPRQLVFRP | 18 | in-silico prediction |
| HEV-ORF1-Y$_{241-256}$ | ORF 1 | Y Domain | 241 | 256 | RTTKIVGDHPLVIERV | 16 | in-silico prediction |
| HEV-ORF1-Y$_{260-275}$ | ORF 1 | Y Domain | 260 | 275 | GCHFVLLLTAAPEPSP | 16 | in-silico prediction |
| HEV-ORF1-Y$_{339-354}$ | ORF 1 | Y Domain | 339 | 354 | RLMTYLRGISYKVTVG | 16 | in-silico prediction |
| HEV-ORF1-Y$_{403-417}$ | ORF 1 | Y Domain | 403 | 417 | AQKFITRLYSWLFEK | 15 | in-silico prediction |
| HEV-ORF1-Y$_{426-440}$ | ORF 1 | Y Domain | 426 | 440 | RQLQFYAQCRRWLSA | 15 | in-silico prediction |
| HEV-ORF1-Cys$_{439-453}$ | ORF 1 | Cysteine protease | 439 | 453 | SAGFHLDPRVLVFDE | 15 | in-silico prediction |
| HEV-ORF1-Cys$_{526-543}$ | ORF 1 | Cysteine protease | 526 | 543 | RQLEALYRALNIPHDIAA | 18 | in-silico prediction |
| HEV-ORF1-Hel$_{1045-1062}$ | ORF 1 | Viral RNA Helicase | 1045 | 1062 | PPHLLLLHMQRASSVHLL | 18 | in-silico prediction |
| HEV-ORF1-Hel$_{1169-1184}$ | ORF 1 | Viral RNA Helicase | 1169 | 1184 | ARGLIQSSRAHAIVAL | 16 | in-silico prediction |
| HEV-ORF1-Hel$_{1189-1203}$ | ORF 1 | Viral RNA Helicase | 1189 | 1203 | EKCVILDAPGLLREV | 15 | in-silico prediction |
| HEV-ORF1-RdRp$_{1430-1446}$ | ORF 1 | RNA-dependent RNA polymerase | 1430 | 1446 | AIEKEILALLPPNIFYG | 17 | in-silico prediction |
| HEV-ORF1-RdRp$_{1497-1513}$ | ORF 1 | RNA-dependent RNA polymerase | 1497 | 1513 | QWLIRLYHLVRSAWILQ | 17 | in-silico prediction |
| HEV-ORF1-RdRp$_{1567-1581}$ | ORF 1 | RNA-dependent RNA polymerase | 1567 | 1581 | CSDYRQSRNAAALIA | 15 | in-silico prediction |
| HEV-ORF1-RdRp$_{1637-1653}$ | ORF 1 | RNA-dependent RNA polymerase | 1637 | 1653 | AVCDFLRGLTNVAQVCV | 17 | in-silico prediction |
| HEV-ORF2$_{7-24}$ | ORF 2 | Capsid | 7 | 24 | LLLFFVLLPMLPAPPAGQ | 18 | in-silico prediction |

| Peptide | ORF | Protein | Start | End | Amino acid sequence | Length | Source |
|---|---|---|---|---|---|---|---|
| HEV-ORF2-S130-147 | ORF 2 | Capsid S-Domain | 130 | 147 | AILRRQYNLSTSPLTSSV | 18 | in-silico prediction |
| HEV-ORF2-S150-164 | ORF 2 | Capsid S-Domain | 150 | 164 | GTNLVLYAAPLNPLL | 15 | in-silico prediction |
| HEV-ORF2-S179-193 | ORF 2 | Capsid S-Domain | 179 | 193 | ASNYAQYRVVRATIR | 15 | in-silico prediction |
| HEV-ORF2-S183-198 | ORF 2 | Capsid S-Domain | 183 | 198 | AQYRVVRATIRYRPLV | 16 | in-silico prediction |
| HEV-ORF2-S188-205 | ORF 2 | Capsid S-Domain | 188 | 205 | VRATIRYRPLVPNAVGGY | 18 | in-silico prediction |
| HEV-ORF2-M368-383 | ORF 2 | Capsid M-Domain | 368 | 383 | IALTLFNLADTLLGGL | 16 | in-silico prediction |
| HEV-ORF2-P459-474 | ORF 2 | Capsid P-Domain | 459 | 474 | SRPFSVLRANDVLWLS | 16 | in-silico prediction |
| HEV-ORF2-P516-530 | ORF 2 | Capsid P-Domain | 516 | 530 | WSKVTLDGRPLTTIQ | 15 | in-silico prediction |
| HEV-ORF2-P534-548 | ORF 2 | Capsid P-Domain | 534 | 548 | KTFYVLPLRGKLSFW | 15 | in-silico prediction |
| HEV-ORF2-P567-584 | ORF 2 | Capsid P-Domain | 567 | 584 | DQILIENAAGHRVAISTY | 18 | in-silico prediction |
| Brown et al 1 | ORF 1 | Viral RNA Helicase | 1031 | 1046 | NGRRVVIDEAPSLPP | 15 | Brown et al. |
| Brown et al 2 | ORF 1 | Viral methyltransferase | 37 | 51 | FLSRLQTEILINLMQ | 15 | Brown et al. |
| Brown et al 3 | ORF 2 | Capsid S-Domain | 241 | 255 | ASELVIPSERLHYRN | 15 | Brown et al. |
| Brown et al 4 | ORF 2 | Capsid M-Domain | 405 | 419 | NGEPTVKLYTSVENA | 15 | Brown et al. |
| Brown et al 5 | ORF 2 | Capsid P-Domain | 535 | 549 | SKTFYVLPLRGKLSF | 15 | Brown et al. |
| Brown et al 6 | ORF 2 | Capsid P-Domain | 584 | 599 | YVLPLRGKLSFWEAG | 15 | Brown et al. |
| HEV-ORF31-18 | ORF 3 | Phosphoprotein | 1 | 18 | MNNMFCALPMGSPCALGL | 18 | OLP library |
| HEV-ORF38-25 | ORF 3 | Phosphoprotein | 8 | 25 | LPMGSPCALGLFCCCSSC | 18 | OLP library |
| HEV-ORF315-32 | ORF 3 | Phosphoprotein | 15 | 32 | ALGLFCCCSSCFCLCCPR | 18 | OLP library |
| HEV-ORF322-39 | ORF 3 | Phosphoprotein | 22 | 39 | CSSCFCLCCPRHRPASRL | 18 | OLP library |

| Peptide | ORF | Protein | Start | End | Amino acid sequence | Length | Source |
|---|---|---|---|---|---|---|---|
| HEV-ORF3$_{29-46}$ | ORF 3 | Phosphoprotein | 29 | 46 | CCPRHRPASRLAAVVGGA | 18 | OLP library |
| HEV-ORF3$_{36-53}$ | ORF 3 | Phosphoprotein | 36 | 53 | ASRLAAVVGGAAAVPAVV | 18 | OLP library |
| HEV-ORF3$_{43-60}$ | ORF 3 | Phosphoprotein | 43 | 60 | VGGAAAVPAVVSGVTGLI | 18 | OLP library |
| HEV-ORF3$_{50-67}$ | ORF 3 | Phosphoprotein | 50 | 67 | PAVVSGVTGLILSPSPSP | 18 | OLP library |
| HEV-ORF3$_{57-74}$ | ORF 3 | Phosphoprotein | 57 | 74 | TGLILSPSPSPIFIQPTP | 18 | OLP library |
| HEV-ORF3$_{64-81}$ | ORF 3 | Phosphoprotein | 64 | 81 | SPSPIFIQPTPLPP | 14 | OLP library |
| HEV-ORF3$_{71-88}$ | ORF 3 | Phosphoprotein | 71 | 88 | QPTPLPPMSYRNPGLELA | 18 | OLP library |
| HEV-ORF3$_{78-95}$ | ORF 3 | Phosphoprotein | 78 | 95 | MSYRNPGLELALDSHPAP | 18 | OLP library |
| HEV-ORF3$_{85-102}$ | ORF 3 | Phosphoprotein | 85 | 102 | LELALDSHPAPSAPLGAT | 18 | OLP library |
| HEV-ORF3$_{92-109}$ | ORF 3 | Phosphoprotein | 92 | 109 | HPAPSAPLGATSPSAPPL | 18 | OLP library |
| HEV-ORF3$_{99-116}$ | ORF 3 | Phosphoprotein | 99 | 116 | LGATSPSAPPLSHVVDLP | 18 | OLP library |
| HEV-ORF3$_{106-122}$ | ORF 3 | Phosphoprotein | 106 | 122 | APPLSHVVDLPQLGLRR | 17 | OLP library |
